# Mitochondrial sodium/calcium exchanger NCLX regulates glycolysis in astrocytes, impacting on cognitive performance

**DOI:** 10.1101/2022.09.16.507284

**Authors:** João Victor Cabral-Costa, Carlos Vicente-Gutiérrez, Jesús Agulla, Rebeca Lapresa, John W. Elrod, Ángeles Almeida, Juan P. Bolaños, Alicia J. Kowaltowski

## Abstract

Intracellular Ca^2+^ concentrations are strictly controlled by plasma membrane transporters, the endoplasmic reticulum, and mitochondria, in which Ca^2+^ uptake is mediated by the mitochondrial calcium uniporter complex (MCUc), while efflux occurs mainly through the mitochondrial Na^+^/Ca^2+^ exchanger (NCLX). RNAseq database repository searches led us to identify the *Nclx* transcript as highly enriched in astrocytes when compared to neurons. To assess the role of NCLX in mouse primary culture astrocytes, we inhibited its function both pharmacologically or genetically. This resulted in re-shaping of cytosolic Ca^2+^ signaling and a metabolic shift that increased glycolytic flux and lactate secretion in a Ca^2+^-dependent manner. Interestingly, *in vivo* genetic deletion of NCLX in hippocampal astrocytes improved cognitive performance in behavioral tasks, whereas hippocampal neuron-specific deletion of NCLX impaired cognitive performance. These results unveil a role for NCLX as a novel modulator of astrocytic glucose metabolism, impacting on cognition.

## Introduction

Ca^2+^ is an important second messenger which participates in a plethora of cell signaling pathways and brain functions, including membrane excitability, synaptic transmission, and plasticity (Kawamoto et al., 2012). Conversely, Ca^2+^ homeostasis disruption occurs under pathological conditions such as senescence and neurodegeneration (Cabral-Costa and Kowaltowski, 2020). Intracellular Ca^2+^ concentrations are tightly controlled by plasma membrane transporters (Kawamoto et al., 2012), the endoplasmic reticulum (Arruda and Parlakgül, 2022), and mitochondria (Cabral-Costa and Kowaltowski, 2020).

We recently found that cerebral mitochondrial Ca^2+^ homeostasis is modulated by dietary calorie intake (Amigo et al., 2017), with strong protective effects against neuronal damage by excitotoxicity, a process that involves loss of cellular Ca^2+^ homeostasis (Arundine and Tymianski, 2003). This shows that these organelles, in addition to their canonical function generating most neuronal ATP, are also important regulators of intracellular Ca^2+^ responses, at least under pathological conditions. However, whether mitochondrial Ca^2+^ homeostasis has physiological impacts on the different cell types of the brain is unknown.

Mitochondrial Ca^2+^ uptake and release were first described in the 1960s (DeLuca and Engstrom, 1961; Vasington and Murphy, 1962; Lehninger et al., 1963; Drahota and Lehninger, 1965), but the major molecular components of the mitochondrial Ca^2+^ handling system were only recently identified (Palty et al., 2010; Perocchi et al., 2010; De Stefani et al., 2011; Baughman et al., 2011; Plovanich et al., 2013; Sancak et al., 2013). Ca^2+^ uptake is mediated by the Mitochondrial Calcium Uniporter (MCU) Complex (MCUc), comprised of a tetramer of MCUs, the structural stabilizer Essential MCU Regulator (EMRE), and gating and regulatory subunits Mitochondrial Calcium Uptake Protein (MICU)-1, −2 or −3 (thoroughly reviewed by Feno et al. (2021). Cerebral mitochondrial Ca^2+^ efflux is mostly mediated by a Na^+^/Ca^2+^ exchanger (NCLX, Fig. 1A), which removes Ca^2+^ from the matrix in exchange for Na^+^ from the intermembrane space (Palty et al., 2010; Assali and Sekler, 2021; Serna et al., 2022).

**Figure 1.**
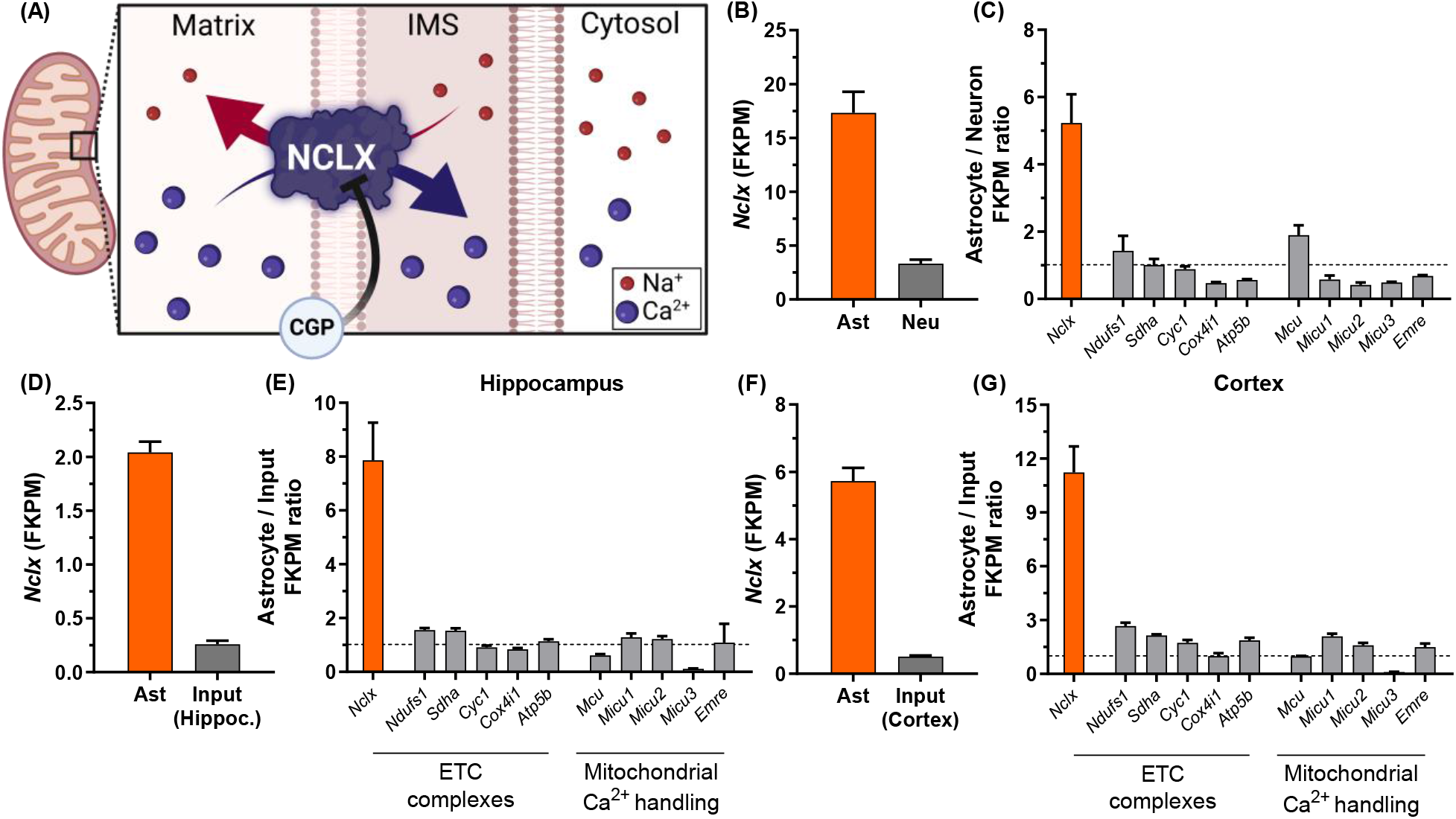
*Nclx* (*Slc8b1, Slc24a6*) transcript is enriched in astrocytes. (A) Schematic illustration of NCLX exchanging extramitochondrial Na^+^ with matrix Ca^2+^, in a manner inhibited by its pharmacological modulator CGP-37157 (CGP). RNA-seq data extracted from Zhang et al. (2014) indicating (B) fragments per kilobase million (FPKM) values of *Nclx* transcript from astrocytes and neurons isolated from mouse cerebral cortex, and (C) FPKM value ratios between astrocytes and neurons from selected transcripts, average ± SD. RNA-seq data extracted from Chai et al. (2017) and Srinivasan et al. (2016) indicating FPKM values for the *Nclx* transcript from isolated astrocytes and respective hippocampal (D) or cortical (F) tissues, and ratio of FPKM values from selected transcripts between astrocytes and input tissue in the hippocampus (E) and (G) cortex. Bars indicate mean ± SEM.

Apart from controlling mitochondrial and cytosolic ion fluxes, NCLX activity has been described to protect hearts against oxidative damage (De La Fuente et al., 2018; Luongo et al., 2017), modulate cardiac hypertrophy (Garbincius et al., 2022), and mediate cellular responses to hypoxia by modulating mitochondrial Na^+^ levels (Hernansanz-Agustín et al., 2020). NCLX also prevents excess intramitochondrial Ca^2+^ in brown adipocytes upon adrenergic activation (Assali et al., 2020), and modulates insulin secretion in β-cells (Nita et al., 2012, 2014, 2015), showing it has important physiological metabolic effects.

In neurons, NCLX integrates mitochondrial metabolism and Ca^2+^ signaling responses (Kostic et al., 2015, 2018), prevents excitotoxicity (Hagenston et al., 2022), and participates in the pathogenesis of forms of Parkinson’s and Alzheimer’s disease (Ludtmann et al., 2019; Jadiya et al., 2019; Britti et al., 2020). Indeed, impaired NCLX activity is associated with reduced synaptic activity and mental retardation (Stavsky et al., 2021). Much less is known about NCLX in astrocytes, although it has been shown that its knockdown impairs proliferation *in vitro* (Parnis et al., 2013) and cell viability *in vivo* (Hagenston et al., 2022).

During search analyses of several public database repositories, we found that NCLX is highly expressed in astrocytes, the most abundant cell types of the brain (Khakh and Deneen, 2019) that participate in neurotransmitter uptake, glutamate recycling, neuronal energy metabolism, and redox balance (Oheim et al., 2018; Bonvento and Bolaños, 2021). Interestingly, we found that, while *in vivo* NCLX deletion in hippocampal astrocytes improves cognitive performance, it leads to cognitive impairment when deleted in hippocampal neurons. These findings correlated with an induction of the glycolytic flux and lactate secretion from astrocytes, revealing that this mitochondrial exchanger has a major impact on brain metabolism and function.

## Results

We were interested in studying the physiological role of mitochondrial Ca^2+^ transport in brain function. Interestingly, there is literature evidence that NCLX, the main mitochondrial Ca^2+^ extrusion pathway (involving exchange for Na^+^ ions, Fig. 1A), is specifically and strongly expressed in astrocytes. To quantify this astrocyte-specific *Nclx* expression, we mined public RNA-seq databases (Chai et al., 2017; Srinivasan et al., 2016; Zhang et al., 2014) and found that *Nclx* mRNA was indeed highly enriched in astrocytes in comparison to neurons (Fig. 1B). This > 5-fold level of enrichment of *Nclx* was a specific astrocytic signature, not associated with total mitochondrial protein, since astrocyte/neuron expression ratios for other mitochondrial proteins, such as those of the electron transport chain and mitochondrial Ca^2+^ transport, were not similarly enriched (Fig. 1C). The enrichment of *Nclx* in astrocytes was also consistent among different databases, and quite significant (8 to 11-fold) when analyzed as astrocyte versus total input tissue in the hippocampus (Fig. 1D,E) and cortex (Fig. 1F,G).

Based on this remarkable enrichment of NCLX specifically in astrocytes, we sought to investigate the effects of this exchanger on astrocytic function. To this end, we used an *in vitro* model of primary murine astrocyte cultures to assess the effects of NCLX inhibition. Parnis et al. (2013) demonstrated that *Nclx* silencing in astrocytes shaped stimulus-induced cytosolic Ca^2+^ responses. In good agreement with this, we observed that pharmacological NCLX inhibition with CGP-37157 (CGP) in cultured astrocytes also modified ATP-induced Ca^2+^ signaling (Fig. 2A,B). CGP-treated astrocytes showed a trend toward smaller ATP-induced Ca^2+^ peaks (Fig. 2C) and increased initial clearance slope (Fig. 2B,D). Indeed, NCLX inhibition significantly decreased the cumulative [Ca^2+^] (area under the curve, AUC, Fig 2E). Our results therefore confirm that NCLX is active in astrocytes, and its activity impacts on cellular Ca^2+^ homeostasis.

**Figure 2.**
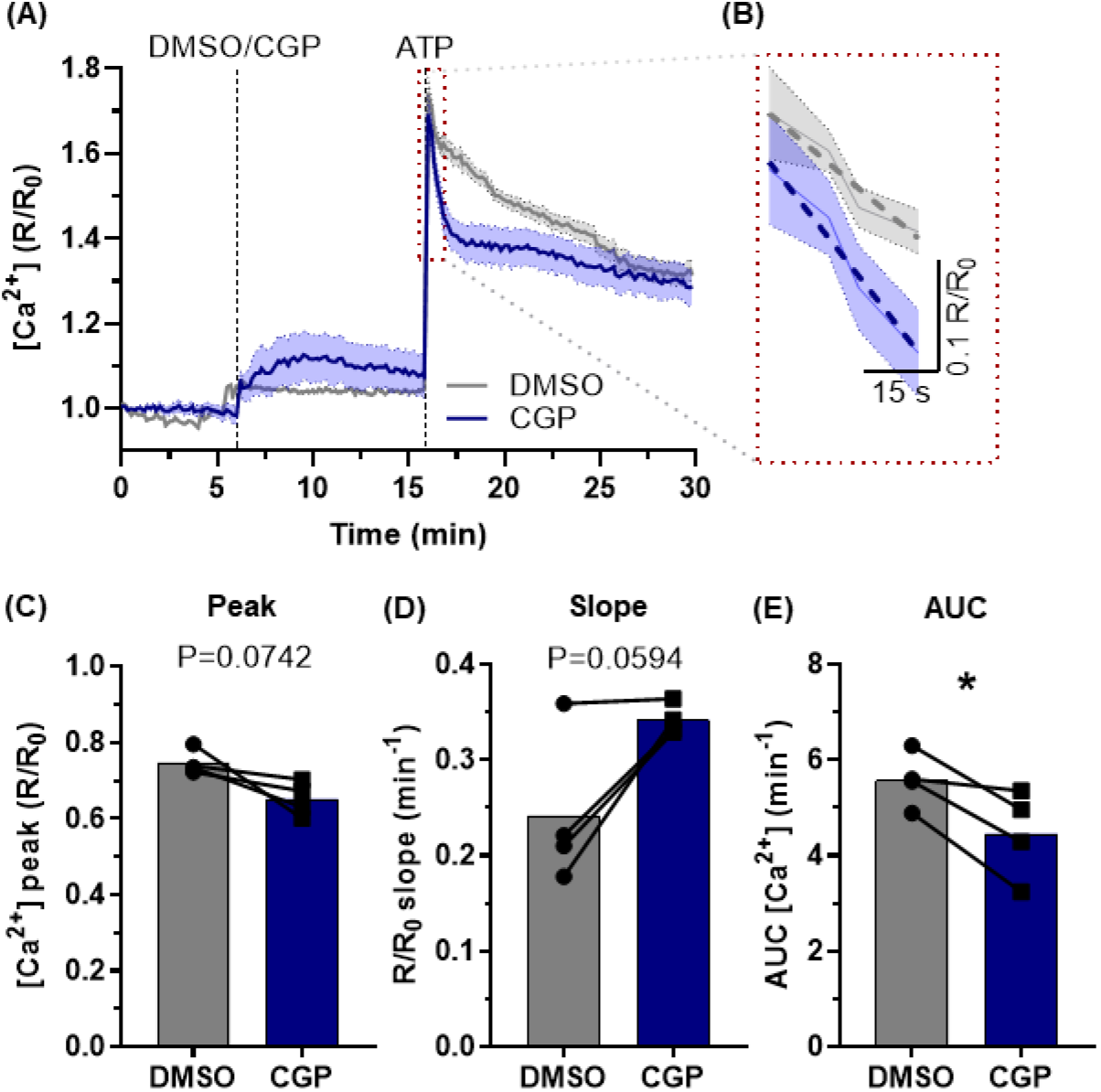
NCLX inhibition changes intracellular Ca^2+^ homeostasis. Primary mouse astrocytes were incubated with the membrane-permeable cytosolic Ca^2+^ probe Fura2-AM and imaged using a fluorescence microscope. (A) Representative trace from a Fura2 imaging experiment (shadowed areas represent the confidence interval; continuous lines indicate mean value from 65 individual cells), indicating incubation with CGP-37157 (or DMSO as control) and ATP to induce a Ca^2+^ wave, and (B) an excerpt highlighting the slope after the peak (dashed lines). (C) Cytosolic [Ca^2+^] peak, (D) slope after reaching peak, and (E) area under the curve (AUC) of the ATP peak. *P < 0.05, paired Student’s t test, n = 4 independent experiments with 55-125 cells each. Paired values are connected by lines, with a bar indicating the mean.

Since NCLX is a mitochondrial protein and modulates intracellular [Ca^2+^], a major metabolic regulator, we next sought to estimate ATP production rates in primary cortical astrocytes acutely stimulated with extracellular ATP with or without NCLX inhibition (Fig. 3), in order to uncover possible metabolic roles for this exchanger. Astrocytic oxygen consumption rates (OCR) and extracellular acidification rates (ECAR) were recorded using a Seahorse Extracellular Flux analysis system, and mitochondrial ATP production and electron transport chain activity were modulated by the addition of oligomycin (an ATP synthase inhibitor) and rotenone plus antimycin A (electron transport inhibitors) (Fig 3A,B). From these traces, the total ATP production rate, as well as its division between oxidative phosphorylation- and glycolysis-associated ATP production rates, were estimated as described by Mookerjee et al. (2017).

**Figure 3.**
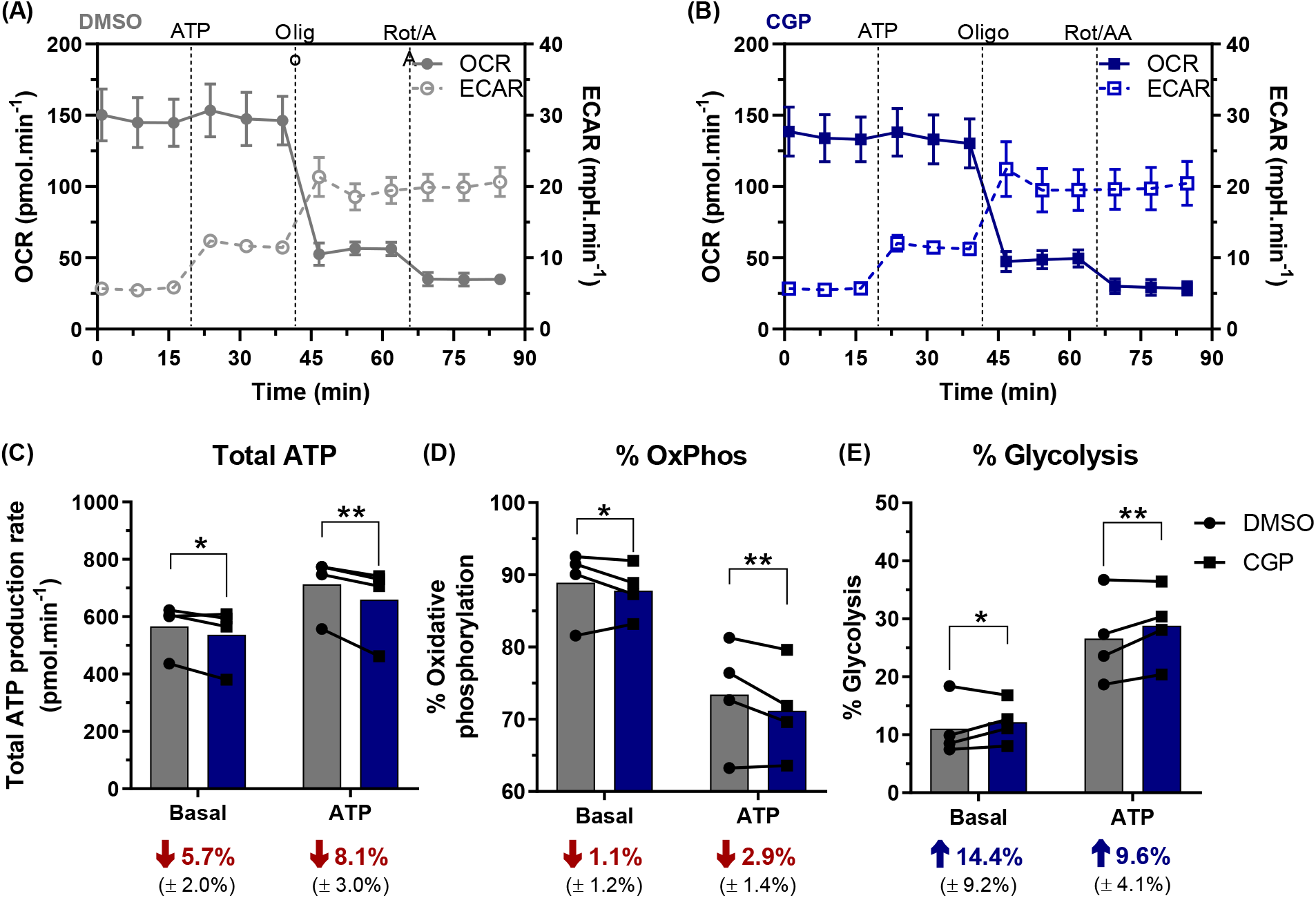
NCLX inhibition increases glycolytic ATP production rates in primary mouse astrocytes. Primary mouse astrocytes incubated with the NCLX inhibitor CGP-37157 (CGP) or DMSO had their oxygen consumption rates (OCR) and extracellular acidification rates (ECAR) monitored in a Seahorse ATP Production Rate assay. Representative traces of (A) DMSO- and (B) CGP-treated astrocytes, stimulated with ATP and followed by oligomycin (oligo) and rotenone plus antimycin A (Rot/AA) inhibition, average ± SEM; Basal and ATP-induced (C) total ATP production rate, and proportional (D) oxidative phosphorylation (OxPhos)- and (E) glycolytic-associated ATP production rate. Average values (± SEM) of the proportional difference between CGP- and DMSO-treated groups were calculated and presented in their respective conditions (C-E). *P < 0.05, **P<0.01, paired 2-way ANOVA followed by Holm-Šidak’s post-hoc test, n = 4 independent experiments. Lines and error bars indicate mean and SD, respectively (A,B); paired values are connected by lines, with a bar indicating the mean (C-E).

NCLX inhibition by CGP-37157 (CGP) induced a decrease in the total ATP production rate (Fig. 3C) both under basal and ATP-stimulated conditions. This was associated with a shift from oxidative phosphorylation to glycolysis (Fig. 3D,E). The increase in glycolysis observed with NCLX inhibition was not exclusive to primary astrocytes. In C6 glioblastoma cells, NCLX inhibition with CGP did not significantly alter overall mitochondrial respiratory parameters (Fig. 3 Suppl 1A-F), but significantly changed ECARs in response to oligomycin (Fig. 3 Suppl 1G), showing a similar metabolic profile to primary astrocytes, which suggests enhanced glycolytic flux. Of note, while the majority of basal ATP production in astrocytes came from mitochondrial respiration (88.9%, Fig. 3D), the effect size of the CGP-induced response was more substantial for glycolytic flux (Fig. 3E), *i.e*. the proportional increase in glycolysis appears to be of greater biological significance than mitochondrial ATP flux reduction.

To further confirm the occurrence of a glycolytic shift promoted by NCLX inhibition, we assessed glucose metabolism through glycolysis by measuring tritiated water (^3^H_2_O) production from radiolabeled glucose, which showed a trend toward an increased glycolytic flux in astrocytes with NCLX inhibition (Fig. 4A). This was paralleled by a significant increase in glucose consumption (Fig. 4B) and lactate secretion (Fig. 4C) under the same conditions, thus confirming that pharmacological inhibition of NCLX activity increases glycolysis and, ultimately, culminates in augmented lactate secretion. This same effect was observed in C6 cells, which presented increased lactate secretion when NCLX was inhibited (Fig. 3 Suppl. 1H).

**Figure 4.**
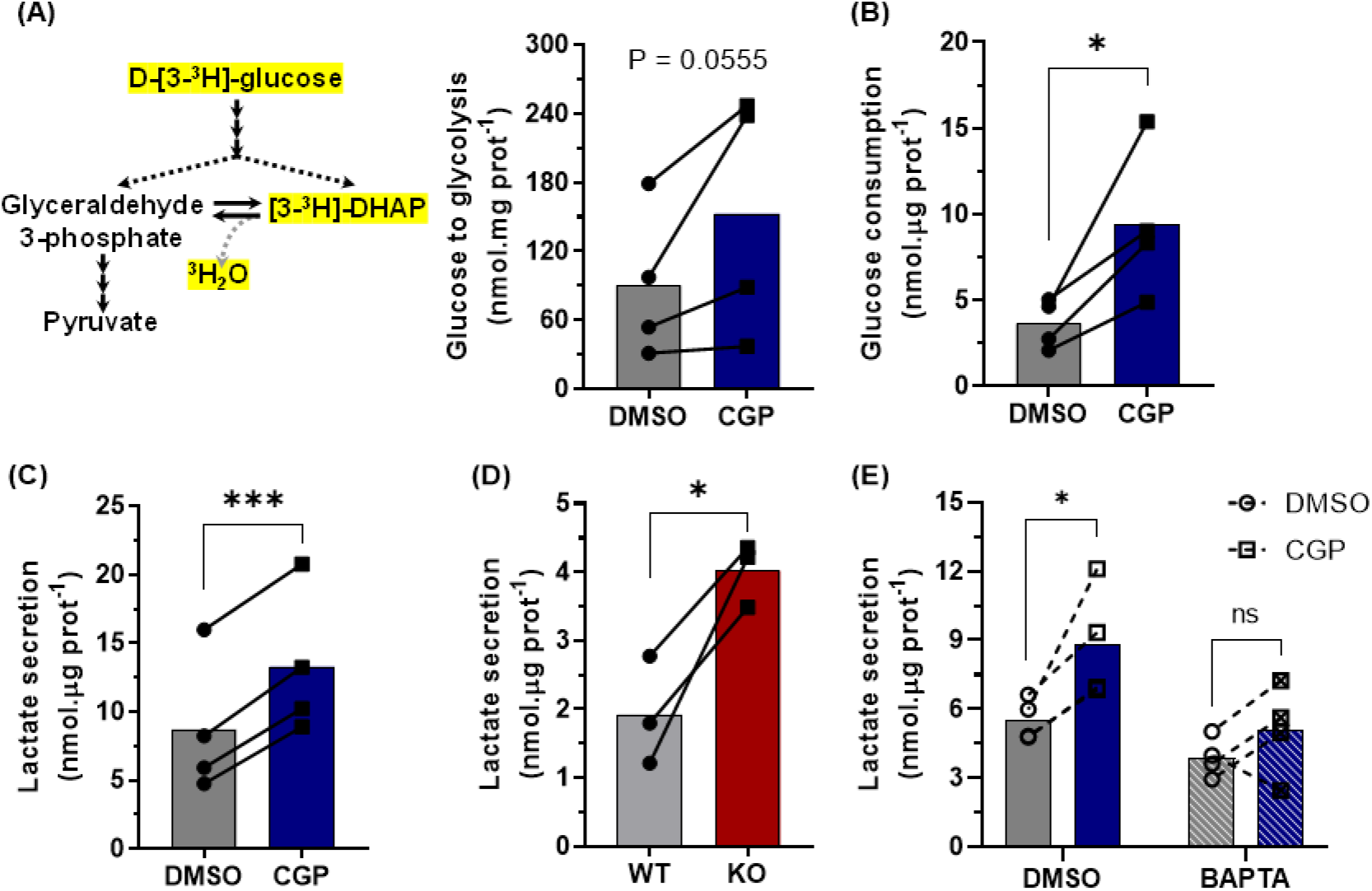
NCLX inhibition increases astrocytic glycolytic flux in a Ca^2+^-dependent manner. Primary mouse astrocytes were co-incubated with the NCLX inhibitor CGP-37157 (CGP) and marked D-[3-^3^H]-glucose for 4 h. Derived tritiated water was measured to estimate glucose metabolism through glycolysis. Glucose consumption (B) and lactate secretion (C) were assessed in parallel experiments. (D) Primary astrocytes derived from *Nclx^loxP/lox^* mice were transduced with an adenoviral vector to express Cre-recombinase and achieve genetic deletion (NCLX KO); lactate secretion was measured during 1 h. Astrocytes were treated with the cytosolic Ca^2+^ chelator BAPTA-AM, followed by incubation with CGP-37157 or DMSO as a control, similarly to Fig 3C. Lactate secretion (D) was then assessed. *P < 0.05, ***P<0.001, paired (B-D) or ratio-paired (A) Student’s t test, or paired 2-way ANOVA followed by Holm-Šidak’s post-hoc test (E), n = 3-4 independent experiments. Paired values are connected by lines, with a bar indicating the mean.

Pharmacological modulations, however, may be prone to undesired off-target effects. We therefore performed experiments in primary cultured astrocytes from *Nclx^loxP/loxP^* mice and induced *Nclx* deletion *in vitro* through adenoviral-mediated Cre expression. While CGP effects are acute (4 h incubations), NCLX knockout was achieved over the course of days, which could lead to compensatory mechanisms and dynamic changes in metabolic modulations observed. Notwithstanding, NCLX knockout induced a significant increase in lactate secretion during 1 h measurements (Fig. 4D), of similar magnitude to those observed in CGP-treated astrocytes.

Since the increased lactate production induced by NCLX inhibition does not involve significantly hampered oxidative phosphorylation or increased ATP demand (Fig. 3), we hypothesized it occurred secondarily to changes in cytosolic Ca^2+^ handling. To investigate this possibility, we verified the effects of NCLX inhibition in astrocytes pre-incubated with the cytosolic Ca^2+^ chelator BAPTA-AM. Again, NCLX inhibition induced a significant increase in lactate secretion in control cells but not in cells in which cytosolic Ca^2+^ was previously chelated by BAPTA (Fig. 4E), thus indicating that Ca^2+^ is necessary for this NCLX-mediated glycolysis modulation. Hence, glycolytic intensification and lactate secretion by NCLX inhibition is a specific, Ca^2+^-dependent effect.

These findings suggest that NCLX has a key functional role in astrocytic metabolic homeostasis, regulating glycolytic flux and lactate secretion. As lactate is secreted from these cells and used as a substrate by neurons, with known effects on memory and synaptic plasticity (Suzuki et al., 2011; Yang et al., 2014; Roumes et al., 2021), we investigated the impact of these metabolic changes on brain function by promoting *Nclx* deletion *in vivo*. Adeno-associated viral vectors were stereotaxically delivered to the hippocampi of *Nclx^loxP/loxP^* adult mice (Fig. 5A) to selectively induce Cre recombinase expression in astrocytes or neurons (Fig. 5B). Behavioral assessment of these animals indicated that neither neuronal nor astrocytic *Nclx* deletion changed their exploratory profile (Fig. 5C,D; Fig. 5 Suppl. 1,2). Surprisingly, astrocytic NCLX KO animals showed improved novel object recognition performance (Fig. 5E; Fig. 5 Suppl. 4) and a similar trend in the Y-maze test (Fig. 5G; Fig. 5 Suppl. 6). In contrast, neuronal *Nclx* deletion negatively influenced the novel object recognition performance (Fig. 5F) without affecting the Y-maze test (Fig. 5H), a result compatible with previous results indicating that increased mitochondrial Ca^2+^ in neurons secondary to NCLX defects is linked to cognitive impairment (Jadiya et al., 2019; Stavsky et al., 2021). These results demonstrate that astrocytic NCLX activity influences cerebral function in a manner associated with enhanced glycolysis and lactate secretion by astrocytes.

**Figure 5.**
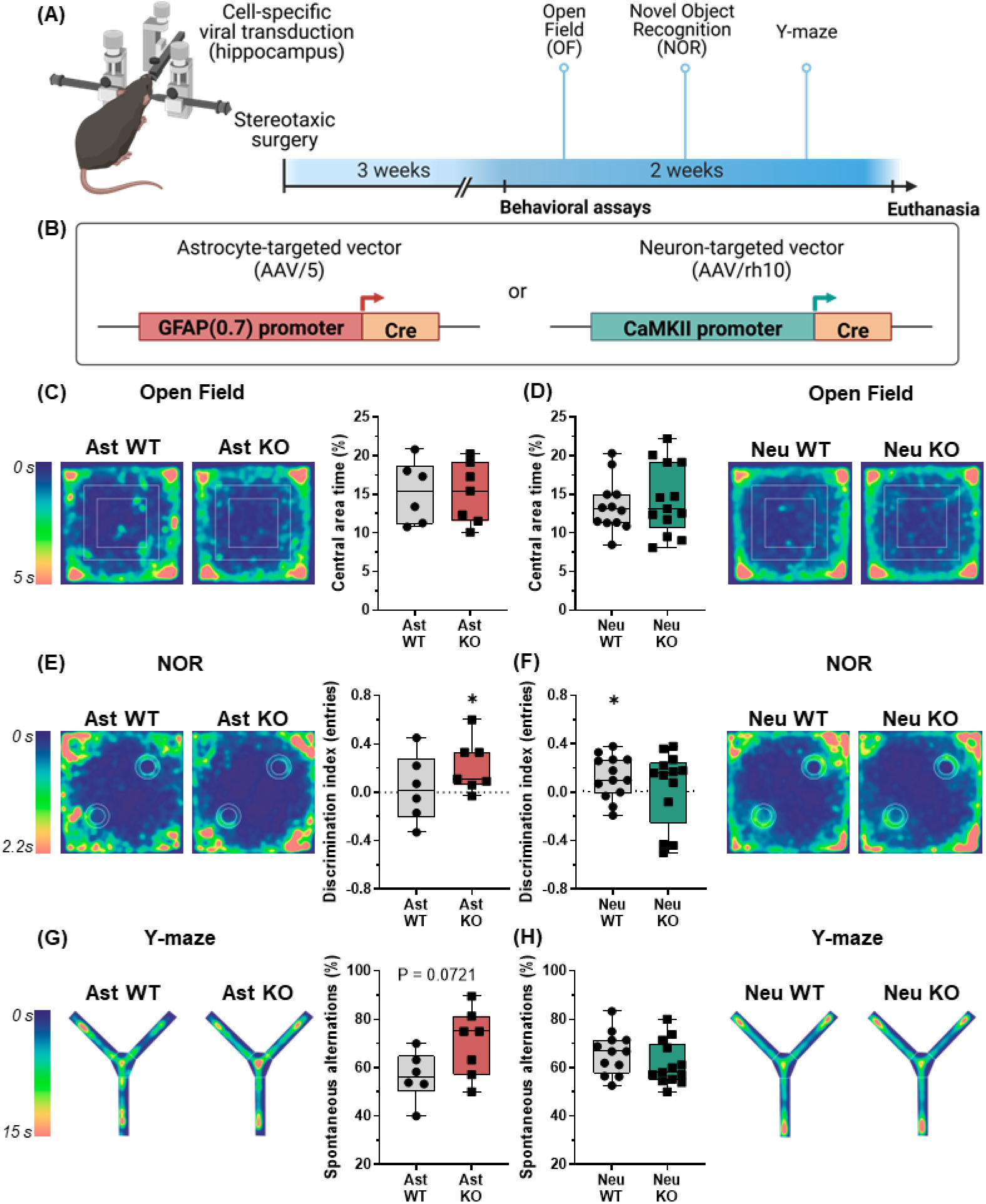
*In vivo* cell-specific NCLX deletion in astrocytes or neurons has opposite behavioral effects. (A) Schematic depiction of the experimental design: *Nclx^loxP/loxP^* mice were injected with cell-targeted vectors stereotaxically in the hippocampus to induce astrocytic or neuronal NCLX deletion, followed by behavioral assessment. (B) Illustration of the viral constructs used to induce astrocyte- (AAV/5) or neuron-specific (AAV/rh10) Cre recombinase expression. (C,D) Open field spatiotemporal quantitative heatmaps showing average occupation of the arena area, and calculation of proportional time in the central area. Not significant, unpaired Student’s t test. (E,F) Novel object recognition spatiotemporal quantitative heatmaps, showing average occupation in the arena during the recognition test, and the discrimination index calculated from entries in novel and familiar object areas. *P < 0.05, one sample Wilcoxon test with theoretical mean = 0.0. (G,H) Y-maze spatiotemporal quantitative heatmaps, showing average occupation of the arena, and calculation of the proportion of spontaneous alternations in respect to total entries. Not significant, unpaired Student’s t test, n = 6-13 mice. Boxes indicate upper and lower quartiles and the median (line), whiskers represent min and max values.

## Discussion

NCLX, the Na^+^/Ca^2+^ exchanger that promotes Ca^2+^ extrusion from mitochondria to the cytosol (Assali and Sekler, 2021; Serna et al., 2022), is highly enriched in astrocytes when compared to other cells in the brain or other mitochondrial proteins (Fig.1; Hagenston et al., 2022). Prior work in astrocytes demonstrated that NCLX silencing leads to impaired astrocyte proliferation *in vitro* (Parnis et al. 2013) and decreased astrocyte numbers *in vivo* (Hagenston et al. 2022). However, little was known about the influence of astrocytic NCLX activity on astrocytic function. NCLX activity in other cell types results mostly in changes in intra and extramitochondrial Na^+^ and Ca^2+^ levels, in a manner dependent on mitochondrial inner membrane potentials (Assali and Sekler, 2021). Indeed, we find that inhibiting NCLX activity significantly impacts on Ca^2+^ homeostasis in astrocytes (Fig. 2).

We also evaluated the effects of NCLX on astrocyte metabolic fluxes, given the known impact of mitochondrial ion transport and Ca^2+^ on metabolic regulation (Llorente-Folch et al., 2015; Juaristi et al., 2019; Ashrafi et al., 2020; Groten and MacVicar, 2022). Interestingly, we found that astrocytic ATP production through oxidative phosphorylation was only slightly decreased by inhibition of NCLX (Fig. 3). Accordingly, Hernansanz-Agustín et al. (2020) did not observe any effect of NCLX activity on mitochondrial respiration in endothelial cells. Furthermore, human colorectal cancer cells present lower maximal respiration in the absence of NCLX, but ATP-linked respiration is unaltered (Pathak et al., 2020). These mild effects of NCLX inhibition on mitochondrial respiration may be related to inhibition of metabolic shuttles secondarily to changes in cytosolic Ca^2+^, since the mitochondrial isoform of glycerol-3-phosphate dehydrogenase (Gherardi et al., 2020) and the aspartate-glutamate exchanger, a component of the malate-aspartate shuttle, are both Ca^2+^-sensitive; the latter is of great relevance for metabolic control in the brain (Llorente-Folch et al., 2015). Hampering the activity of these critical points for mitochondrial NADH uptake decreases maximal electron transport capacity in mitochondria and may also lead to enhanced cytosolic NADH levels (Wang et al., 2022) (Fig. 6). Consistently, accumulation of Ca^2+^ in cerebral mitochondria leads to accumulation of NADH (Díaz-García et al., 2021).

**Figure 6.**
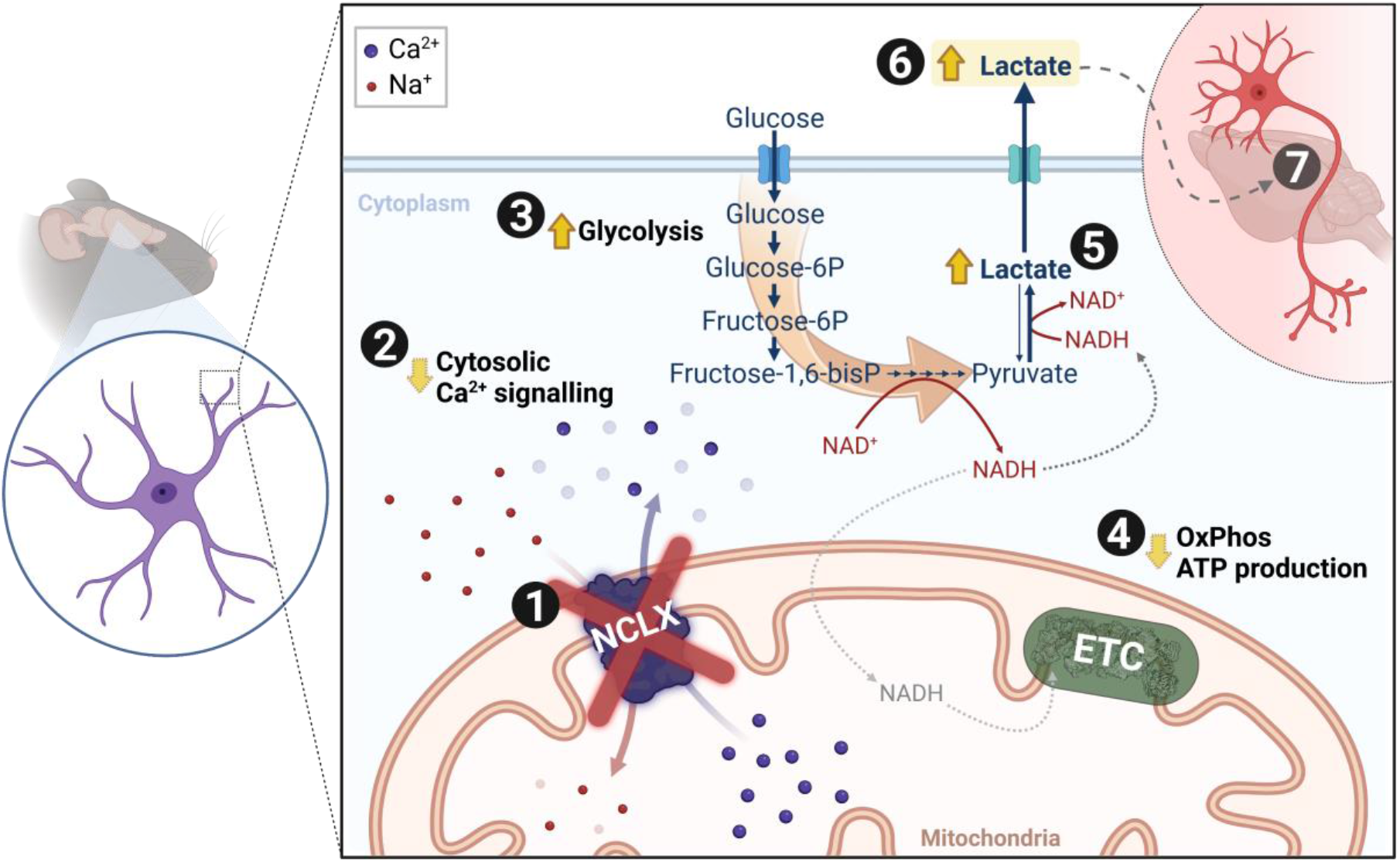
Schematic overview. In astrocytes, (1) inhibition/deletion of mitochondrial Na^+^/Ca^2+^ exchanger (NCLX) activity leads to (2) augmented cytosolic Ca^2+^ clearance. This results in (3) increased glycolytic flux; and (4) slightly decreased mitochondrial oxidative phosphorylation, leading to (5) increased lactate dehydrogenase (LDH)-mediated reduction of pyruvate to lactate. The resulting increased lactate in astrocytes (6) is secreted (7) and may contribute to enhanced behavioral performance *in vivo*. (ETC: electron transport chain).

While the effects of astrocyte NCLX inhibition on mitochondrial electron transport were small, glycolytic ATP fluxes were substantially increased, both in cells with pharmacologically-inhibited NCLX and in knockout cells (Figs. 3,4). Loss of NCLX activity in colorectal cells was also found to significantly increase glycolytic flux. Interestingly, NCLX is modulated by PKA (Assali et al., 2020; Kostic et al., 2015; Zhou et al., 2021), an important metabolic regulatory hub that also influences glycolysis (Rider et al., 2004), further supporting a role for this transporter in the regulatory network of glycolytic activity. Increased glycolytic flux, especially when in the presence of decreased oxidative phosphorylation and lower NADH oxidation (Rigoulet et al., 2020), typically promotes enhanced lactate production. Indeed, we find that astrocytes and C6 glioma cells secrete more lactate when NCLX is inhibited pharmacologically or knocked out (Fig. 5). This effect is a result of changes in cytosolic Ca^2+^ signaling promoted by NCLX, as it was abrogated by the presence of BAPTA as a cytosolic Ca^2+^ chelator.

Astrocytic lactate has long been characterized as a fundamental substrate for neurons (Pellerin and Magistretti, 1994; Herrero-Mendez et al., 2009; Rodriguez-Rodriguez et al., 2012; Mächler et al., 2016; Bonvento and Bolaños, 2021), which also acts as a gliotransmitter, promoting synaptic plasticity, and higher functions (Suzuki et al., 2011; Yang et al., 2014; Adamsky et al., 2018; Jimenez-Blasco et al., 2020; Roumes et al., 2021; Akther and Hirase, 2022). Our data show that NCLX can control lactate secretion and therefore potentially act as a modulator of the astrocyte-to-neuron lactate shuttle, by acting as a connection between cytosolic and mitochondrial Ca^2+^ signaling and glycolytic flux. Indeed, we observed that astrocyte-specific NCLX deletion in the hippocampus improves aspects of mouse cognitive performance (Fig. 5E,G), while hampering NCLX activity in neurons promotes deleterious effects (Kostic et al., 2015; Sharma et al., 2017; Jadiya et al., 2019; Stavsky et al., 2021; Britti et al., 2020, 2021; Hagenston et al., 2022).

In conclusion, we demonstrate that NCLX, which is over-enriched in astrocytes, modulates astrocytic glycolytic flux and lactate secretion secondarily to shaping cytosolic Ca^2+^ signaling (Fig. 6). By fine-tuning astrocytic glycolysis and lactate secretion, NCLX may act as a control check point in brain metabolism impacting on the astrocyte-to-neuron lactate shuttle and cerebral function.

## Materials and methods

### RNAseq public databases

RNAseq data was mined from the public databases published by Zhang et al. (2014), accessed at https://www.brainrnaseq.org/ (last access: 2021-11-01), GEO accession number GSE52564; and by Chai et al. (2017) and Srinivasan et al. (2016), accessed at http://astrocyternaseq.org/ (last access: 2021-11-01), GEO accession numbers GSE84540 and GSE94010, respectively.

### Animal care

Experimental design and animal care standards followed ARRIVE 2.0 guidelines (Percie du Sert et al., 2020). Animal procedures were performed according to Protocol #82/2017 from the *Comissão de Ética em Cuidado e Uso Animal do Instituto de Química da Universidade de São Paulo* and by the Bioethics Committee of the University of Salamanca (reference 449), following requirements described by the *Sociedade Brasileira de Ciência de Animais de Laboratório*, European Union Directive 86/609/EEC and Recommendation 2007/526/ EC, regarding the protection of animals used for experimental and other scientific purposes, and enforced under Spanish legislation directive RD1201/2005. Adult mice were maintained in groups of 4-5 animals per cage at the specific pathogen free Animal Experimentation Facility of the University of Salamanca (Biosafety Level 2 environments). Neonates (0-1 days-old) were obtained from breeding cages (1 male and 1-3 females per cage) from the specific pathogen free Animal Care Facility of the Institute of Chemistry and Faculty of Pharmaceutical Sciences at the University of São Paulo and from the Animal Experimentation Facility of the University of Salamanca. All animals were maintained in a light-dark cycle of 12 h, 45-65% humidity, 20-25 °C, with open and unlimited access to standard solid diet and water, in a microisolator system. Cages were changed and sanitized 1-2 times/week.

The number of neonates was determined by the demand of primary astrocyte cultures. Protocols and study design were optimized to yield the maximal cell count using the smallest numbers of animals, in accordance with the 3Rs principle (Percie du Sert et al., 2020). For *in vivo* experiments, a limited sample size was allocated, as the initial objective was to conduct an exploratory assessment in pursuit of evidence pointing out effects that may be of interest for further investigation. Experimental feasibility (surgery, recovery, behavioral assays, euthanasia) and operational limitations (processing capacity, total study duration and budget) were taken into account and adjusted as in a Fermi’s approximation (Reynolds, 2019). Sample size range is specified in each figure legend and every animal is depicted as a symbol in graphical representations. Animals were allocated to each group haphazardly and evenly through experimental and control groups, and cage order was counterbalanced through the course of experimental assays to avoid a time of the day bias. In total, 43 adult mice were used for *in vivo* experiments, 3 of which were excluded due to surgery issues.

*In vitro* pharmacological experiments with primary astrocytes were conducted in cells from C57Bl/6NTac mice. *Nclx^loxP/loxP^* (originally denoted *Slc8b1^fl/fl^*) mice were designed and produced at Dr. John Elrod’s lab, as described by Luongo et al. (2017). Parental breeding pairs were kindly provided and shipped by Dr. Antonio Martínez-Ruiz (Hospital Universitario de La Princesa, Madrid, Spain) and maintained using a C57Bl/6J background.

### Cell cultures

Mouse cortical astrocyte primary cultures were conducted as previously described (Jimenez-Blasco et al., 2020). Briefly, brains of neonates (P0-1, both male and female) were dissected and digested with 0.1% trypsin (#T0134, Sigma-Aldrich, Saint Louis, MO, USA) in the presence of 60 μg/mL DNAse I (#DN25, Sigma-Aldrich) in HBSS medium (#14175095, Gibco, Life Technologies, Carlsbad, CA, USA). The tissue was then dissociated in HBSS containing 24 μg/mL DNAse I, decanted, and the resulting cell suspension was counted, plated, and maintained in Low Glucose DMEM (5.5 mM glucose, 1 mM pyruvate, 4 mM glutamine; #31600034, Gibco) supplemented with 10% fetal bovine serum (#12605729, Gibco) and 1% penicillin/streptomycin (#15140122, Gibco), in a 5% CO_2_, 37°C, humidified incubator. Cells were grown in a 75 cm^2^ flask for 7 days and then shaken at 200 rpm in an incubator; the supernatant was discarded, and the remaining astrocyte-enriched culture was re-seeded at 50.10^3^/cm^2^ and grown for 3-7 days for the experiments.

The C6 cell line stock (BCRJ Cat# 0057, RRID:CVCL_0194) was kindly donated by Dr. Cristoforo Scavone (Institute of Biomedical Sciences, University of São Paulo, São Paulo, Brazil). C6 cells were grown and maintained in High Glucose DMEM (25 mM glucose, 1 mM pyruvate, 4 mM glutamine; #12800017, Gibco) supplemented with 10% fetal bovine serum and 1% penicillin/streptomycin, in a 5% CO_2_, 37°C, humidified incubator.

For the experiments, unless otherwise stated, all cell media were changed for a respective serum-free version and cells were allowed a 1 h equilibration period, after which 10 μM CGP-37157 (#1114, Tocris, Bio-Techne, Bristol, UK) or sterile DMSO, as a control, were added. When necessary, BAPTA-AM (10 μM, #A1076, Sigma) or DMSO, as a control, was incubated over the last 30 min of the equilibration period.

### Seahorse assays

Purified astrocytes or C6 cells were plated at a density of 30·10^3^ or 72·10^3^ cells per well, respectively, on XFe24 Seahorse plates (#100777-004, Agilent, Santa Clara, CA, USA) and experiments were conducted at day *in vitro* (DIV) 15 ± 1, either for acute pharmacological inhibition of NCLX or for NCLX knock-out (7 days after viral transduction). Cells were washed once with experimental medium – DMEM (phenol-free, lacking sodium bicarbonate; #D5030, Sigma-Aldrich) supplemented with 1 mM pyruvate, 4 mM glutamine, 10 mM HEPES, 1% penicillin/streptomycin, and 5.5 mM glucose (for astrocytes) or 25 mM glucose (for C6 cells) – and pre-incubated at 37°C, room atmosphere, for 1 h in 500 μL experimental medium. Tests were conducted as described below, using pre-titrated inhibitor concentrations, and assessing respective oxygen consumption rates (OCR) and extracellular acidification rates (ECAR).

Astrocyte experiments were normalized by automated cell count, as described previously (Assali et al., 2020). Seahorse XFe24 plates were washed once with PBS right after ending the assay and fixed overnight at 4°C with PFA 4% in methanol, DAPI-stained and imaged and analyzed in a custom workflow on a High Content Screening Operetta CLS apparatus (Perkin Elmer, MA, USA). Alternatively, C6 cells were normalized by determination of total protein concentration through a BCA kit (Thermo Fisher Scientific, Rockford, USA).

#### ATP rate test

Total ATP production rates were estimated as previously described (Kakimoto et al., 2021; Mookerjee et al., 2017). Astrocytes were pre-incubated with 10 μM CGP-37157 or DMSO for 1 h and then plates were inserted in a XFe24 Seahorse Analyzer apparatus (#102238-100, Agilent). Cells were stimulated with 100 μM ATP, followed by ATP synthase inhibition with oligomycin (oligo, 2.5 μM) and electron transport chain inhibition with rotenone (rot, 1.0 μM) plus antimycin A (AA, 2.0 μM). Total ATP production rates, as well as its partition between glycolytic and oxidative phosphorylation, were calculated according to the manufacturer’s instructions (Romero et al., 2018), considering standard values of required constants and the buffer factor as 3.13 mM/pH.

#### MitoStress test

C6 cell metabolic assessment was conducted using a MitoStress test, as previously described (Amigo et al., 2017). Cell plates were inserted in an XFe24 Seahorse Analyzer apparatus (#102238-100, Agilent), acutely challenged with 10 μM CGP-37157 or DMSO, followed by ATP synthase inhibition with oligomycin (oligo, 0.5 μM), mitochondrial uncoupling with 2,4-dinitrophenol (DNP, 200 μM), and electron transport chain inhibition with rotenone (rot, 1 μM) plus antimycin A (AA, 1 μM). Non-mitochondrial respiration is defined as the Rot+AA-insensitive OCR and is subtracted from other parameters; OCR_CGP_ was derived from the average between the last 3 OCR measurements; OCR_proton-leak_ was calculated from the average between the last two oligomycin-insensitive OCR measurements; OCR_ATP-linked_ was calculated from the difference between OCR_CGP_ and OCR_proton-leak_; ΔECAR_oligo_ was calculated as the difference between the first ECAR measurement right after and the one right before the oligomycin addition.

### Glycolytic flux

Glucose-to-glycolysis metabolism was assessed as described elsewhere (Jimenez-Blasco et al., 2020). In brief, astrocytes were washed and maintained in experimental medium at 37°C, room atmosphere, for 1 h to equilibrate. Then, cells were incubated with [3-^3^H]-glucose (2 μCi/well) and 10 μM CGP-37157 or DMSO for 4 h, under gentle orbital rotation (60 rpm) at 37°C. Reactions were stopped by acidification with 20% perchloric acid, and cell media was collected and allowed to equilibrate with a separated tube containing 500 μL deionized water, enclosed in a sealed glass vial, and maintained in a rotating incubator at 60 rpm, 37°C, room atmosphere, for 72 h. Produced ^3^H_2_O was indirectly measured from these plastic vials through liquid scintillation counting (Tri-Carb 4810 TR, PerkinElmer).

### Lactate secretion and glucose consumption

In brief, cells were washed and maintained in serum-free culture medium for 1 h for equilibration, collected (baseline measurement), and followed by incubation with 10 μM CGP-37157 or DMSO. Cell medium was collected right after CGP addition and after 1 or 4 h, and both glucose and lactate (Vicente-Gutierrez et al., 2019) levels were measured spectrophotometrically. Lactate concentrations were determined through assessment of NADH formation at λ = 340 nm in a buffer (250 mM glycine, 500 mM hydrazine, 1 mM EDTA, pH 9.5) containing 1 mM NAD^+^ and 22.5 U/mL lactate dehydrogenase, or using a commercial kit (#138, Labtest, Lagoa Santa, MG, Brazil). Glucose concentrations were determined by following NADPH production at λ = 340 nm in a tris buffer (100 mM, pH 8.0), containing 0.5 mM MgCl2, 2 mM ATP, 1.5 mM NADP^+^, 2.5 U/mL hexokinase and 1.25 U/mL glucose-6-phosphate dehydrogenase.

### Viral transduction

Cre recombinase expression was induced *in vitro* through an adenoviral vector (Ad5-CMV-Cre-eGFP, lot# Ad4334 13D6, University of Iowa Viral Vector Core, Iowa City, IA, USA) or its respective empty vector as a control (Ad5-CMV-GFP, lot# Ad4415 13D3, University of Iowa Viral Vector Core). Primary astrocytes were infected 2 days after being re-plated at 15 MOI (multiplicity of infection). Virus suspension was removed 24 h after transduction and experiments were conducted 7 days after beginning of infection.

For *in vivo* experiments (Fig. 5B), Cre recombinase expression was mediated by adeno-associated viral vectors (AAV, all from Vector Biolabs, Malvern, PA, USA) and driven by an astrocyte-specific GFAP promoter (AAV/5-GFAP(0.7)-GFP-2A-iCre, #VB1131, lot# 190527#25) or by a neuronal-specific CaMKII promoter (AAV/rh10-CamKII(0.4)-eGFP-T2A-Cre, #VB1435, lot# 201123#1). Control groups were transduced with the empty vectors AAV/5-GFAP(0.7)-eGFP (#VB1149, lot# 190527#24) and AAV/rh10-CamKII(0.4)-eGFP (#VB1435, lot# 201123#1), respectively.

### Stereotaxic surgery

Surgery was conducted as described by Lapresa et al. (2022). Male *Nclx^loxP/loxP^* mice (11.7 ± 2.5 weeks old) were briefly anesthetized with sevoflurane (4% for induction, 2.5% for maintenance) in a 30% O_2_ and 70% N_2_O atmosphere (0.4 and 0.8 L/min, respectively). Animals were appropriately positioned in the stereotaxic apparatus (#1900, David Kopf Instruments, Tujunga, CA, USA) coupled with a digital readout (Wizard 550, Anilam, ACU-RITE/Heidenhain Corporation, Schaumburg, IL, USA), maintained under a heat lamp, and had their temperatures monitored by a rectal thermometer. Injections were controlled by a digitally-controlled pump (UltraMicroPump with a Micro4 UMC4 III controller, World Precision Instruments, Sarasota, FL, USA), in which 2 μl containing 1 . 10^10^ PFU/μL of AAV/5 vectors (for astrocytic deletion, see constructs above) or 2.75 . 10^12^ viral genome copies/μL of AAV/rh10 vectors (for neuronal deletion, see constructs above) diluted in sterile PBS with 0.001% Pluronic F-68 were administered bilaterally in two depths (1 μl each) at 500 nL/min. Hippocampi were targeted according to the following coordinates, based on Paxinos and Franklin atlas (Paxinos and Franklin, 2001) and previously validated (Jimenez-Blasco et al., 2020): AP = −2 mm, ML = ± 1.5 mm, and DV = −2 mm (first injection) and −1.5 mm (second injection). Animals were kept in heated cages and closely monitored up to full recovery from anesthesia, and then were observed for the following days. Detection of unexpected recovery issues (*e.g*., infection or excessive inflammation at suture site, inadequate wound healing) or cases when stereotaxic surgery was identified as unsuccessful by the surgeon (*e.g*., syringe content overflowed injection site, death during surgery) were exclusion criteria, and animals were cared for to minimize suffering and euthanized.

### Behavioral assays

Behavioral assessment started 3 weeks after surgery (Fig. 5A), to allow proper recovery and gene recombination. Mice were acclimatized to the experimenter (male researcher) 1 week prior to the beginning of the behavioral assays by daily soft manipulation, and to the experimental room for 1 h before each assay. Assays were tracked by ANY-maze software with the Ami-maze interface in an ANY-box core (40 × 40 cm; Stoelting Co., Wood Dale, IL, USA), except for the Y-maze test, which was conducted on a specific apparatus and manually scored. The experimenter was blinded to animal genotype through all behavioral experiments. Censoring was proceeded when justified by statistical outlier assessment or in the case of operational problems (*e.g*., video recording issue), and properly reported when done.

#### Open Field test

Exploratory behavior was assessed through the Open Field test (Cabral-Costa et al., 2018; Lapresa et al., 2022). Animals were allowed to individually explore the experimental apparatus for 10 min. Total distance, mean speed, time freezing, number of rearings, and central area (defined as a virtual central 20 × 20 cm square) number of entries, and total time were measured.

#### Novel Object Recognition test

On the following day, animals were submitted to 2 sessions of 5 min each, separated by a 30 min interval. These sessions consisted of a training stage (two equal wooden objects on opposite symmetric sides of the arena, Fig. 5E,F) and novel object recognition (NOR, where a second, novel, object substituted one of the familiar ones, at the bottom left position). Total distance, time spent exploring and number of interactions (entries) with each object was measured. NOR discrimination indexes were assessed as an indicator of short-term recognition memory (Cabral-Costa et al., 2018; Vicente-Gutierrez et al., 2019), calculated as the difference in number of interactions (or time) between the novel and the familiar object divided by total number of entries (or total exploration time).

#### Y-maze test

Spontaneous alternation on a Y-maze was assessed as an indicator of working memory (Jimenez-Blasco et al., 2020). Animals were positioned in the central area of a Y-maze, facing the wall on the opposite side of the experimenter, and allowed to explore the maze for 5 min. Entrances on each arm (A, B, C) were manually scored from the recorded video by an independent researcher who was blinded for genotype. Spontaneous alternation was defined as the total number of triads of sequential entrances in three different arms and calculated in Rstudio (version 2022.02.0, PBC, Boston, MA, USA) using the script annotated at https://github.com/jvccosta/NCLXAstMetab.

### Calcium imaging

Calcium levels were live monitored in attached astrocytes through the ratiometric probe Fura-2-AM (#F1221, Invitrogen, Waltham, MA, USA) as done by Kowaltowski et al. (2019). Briefly, cells were plated in glass-bottom culture dishes (#627871, Greiner Bio-One, Kremsmünster, Austria), incubated with 5 μM Fura-2-AM for 30 min at 37°C in experimental medium lacking FBS and supplemented with 1 mg/mL bovine serum albumin (BSA). Fluorescence was assessed at λ_ex_ = 340 (F_340_) and 380 nm (F_380_) and λ_em_ = 510 nm in a Leica DMi-8 microscope equipped with a Fura-2 filter (Leica Microsystems, Buffalo Grove, IL, USA). Cells were followed through additions of CGP (10 μM), ATP (100 μM), as well as ionomycin (20 μM) to allow calibration. Analyses were conducted through FIJI ImageJ 1.52p (Schindelin et al., 2012), in which individual cells (55-125/group per experiment) were identified as regions of interest (ROI) and [Ca^2+^] variation was estimated as the ratio (R) between F_340_/F_380_. Data was calibrated by the maximal ratio induced by ionomycin, controlled for background fluorescence oscillations, and normalized by the initial ratio (R_0_).

### Statistical analyses

All raw data was organized and analyzed in Microsoft Excel (Microsoft 365 MSO, version 2207, Microsoft Corporation, Redmond, WA, USA), and statistical analyses were performed in GraphPad Prism 8 (version 8.4.3, GraphPad Software, San Diego, CA, USA), in which all figures were also plotted. According to the experimental design, as appropriately described in the figure legends, data was analyzed through unpaired, paired, or ratio-paired Student’s t-test; one-sample Wilcoxon test with theoretical mean = 0.0, and paired two-way ANOVA, followed by Holm-Šidak’s post-hoc test for parametric analyses; and through Mann-Whitney test for non-parametric analyses. A ROUT outlier test, with 5% sensitivity, was used to search for outliers.

## Data availability

All raw data will be made available as supplementary material.

## Acknowledgements

We would like to thank Camille C. Caldeira da Silva, Sirlei Mendes de Oliveira, and Monica Resch for the outstanding technical support; the IQ-FCF/USP and IBFG-USAL animal facilities staffs, in the name of Silvania Neves, Renata Spalutto Fontes, Flavia Ong, Monica Carabias-Carrasco, Lucia Martin, and Estefania Prieto-Garcia, for the exceptional animal care; André Costa Oliveira, for contributing with coding guidance; and Amanda Midori Matumoto, for contributing with animal behavior scoring. We are also deeply grateful to Dr. Antonio Martínez-Ruiz for kindly mediating the *Nclx^loxP/loxP^* mouse logistics, as well as to Dr. Pamela Kakimoto, Dr. Bruno Chausse, Dr. José Carlos de Lima Jr., Dr. Marcus F. Oliveira, Dr. Ruben Quintana-Cabrera, Dr. Marcel Vieira-Lara, Dr. Nathalia Dragano, Vitor M. Ramos, Paula Alonso-Batán, Dr. Daniel Jiménez-Blasco, and Dr. Pablo Hernansanz-Agustín for contributing with scientific discussions and inputs. Illustrations were created with Biorender.com. Electron transport chain scheme was constructed based on actual complex crystal structures (PDB #6G2J, 1ZOY, 3CX5, 3HB3, 1CRH, 7TK4).

## Funding

Authors were supported by grant #2020/06970–5 from *Fundação de Amparo à Pesquisa do Estado de São Paulo* (FAPESP); *Centro de Pesquisa, Inovação e Difusão de Processos Redox em Biomedicina* (CEPID Redoxoma, FAPESP grant #2013/07937–8); *Conselho Nacional de Desenvolvimento Científico e Tecnológico* (CNPq); and *Coordenação de Aperfeiçoamento de Pessoal de Nível Superior* (CAPES) line 001; *Agencia Estatal de Investigación* (PID2019-105699RB-I00/AEI/10.13039/501100011033, PDC2021-121013-I00 and RED2018-102576-T to JPB); *Plan Nacional de Drogas* (2020I028 to JPB); *Instituto de Salud Carlos III* (PI21/00727 and RD21/0006/0005 co-funded by the European Union FEDER/FSE+ and NextGenerationEU to AA); and *Junta de Castilla y León* (CS/151P20 co-funded by P.O. FEDER to AA; *Apoyo Regional a la Competitividad Empresarial*, ICE 04/18/LE/0017 to JPB, and *Escalera de Excelencia* CLU-2017-03 to JPB and AA). JVCC was also supported by FAPESP fellowships #2017/14713-0 and #2019/22178-2.

## Conflict of Interest

The authors declare that they have no conflicts of interest with the contents of this article.

## Author contributions

**Conceptualization:** JVCC, JPB, AJK. **Methodology:** JVCC, JWE. **Software:** JVCC. **Validation:** JVCC, CVG. **Investigation:** JVCC, CVG, JA, RL. **Formal Analysis:** JVCC, JA, RL. **Data Curation:** JVCC. **Visualization:** JVCC. **Writing – Original Draft:** JVCC. **Writing – Review:** JA, RL, JWE, AA, JPB, AJK. **Writing – Editing:** JVCC, AA, JPB, AJK. **Resources:** JWE, AA, JPB, AJK. **Funding Acquisition:** JVCC, AA, JPB, AJK. **Supervision:** CVG, JA, AA, JPB, AJK **Project Administration:** JPB, AJK.

## Figure Supplements

**Figure 2 – Supplement 1. DMSO and CGP FURA-2 imaging representative videos.**

**Figure 3 – Supplement 1.**
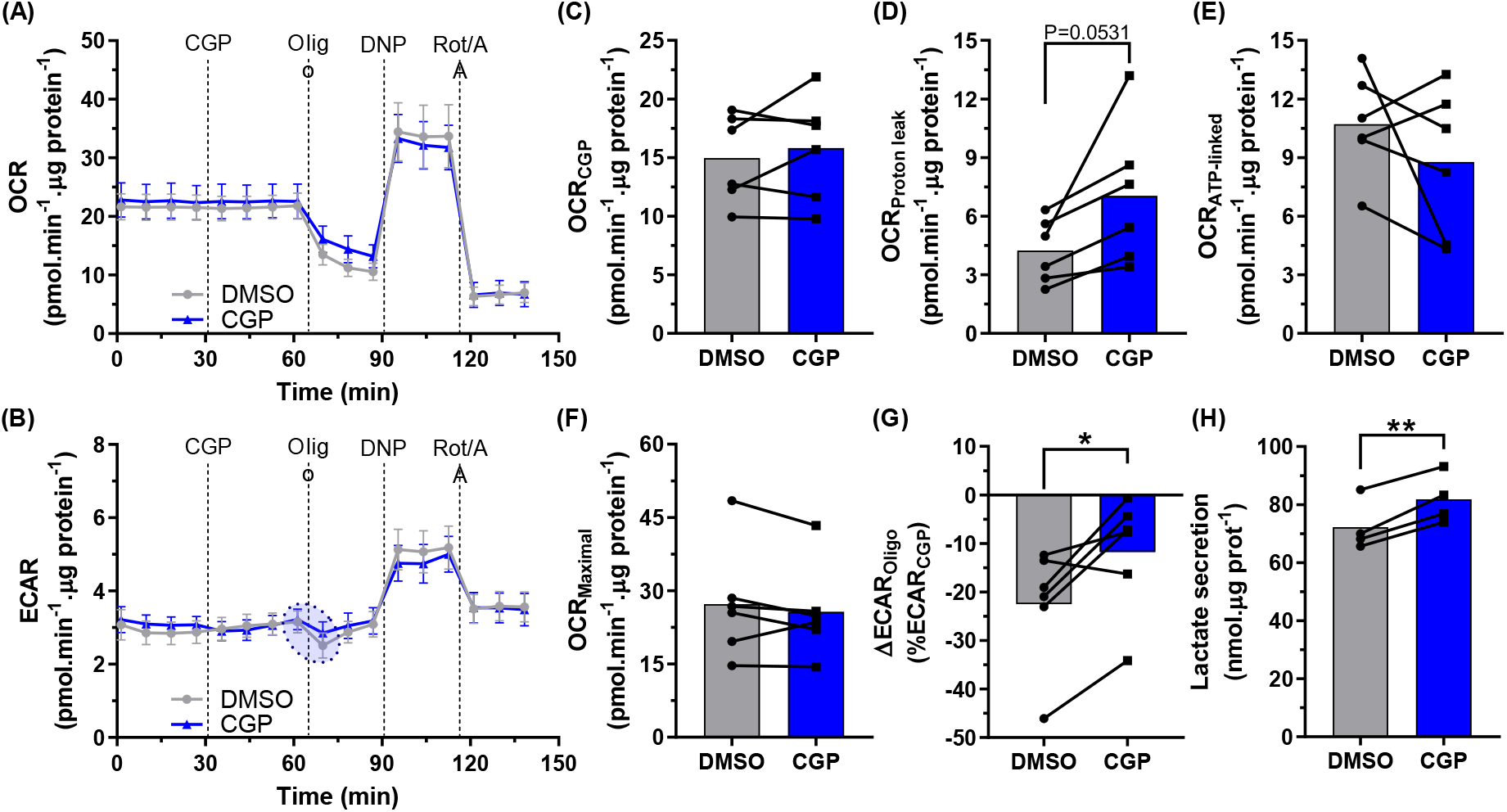
C6 cells present increased lactate secretion upon NCLX inhibition. C6 cells incubated with the NCLX inhibitor CGP-37157 (CGP) or DMSO had their oxygen consumption rate (OCR) and extracellular acidification rate (ECAR) monitored in a MitoStress Seahorse assay. Representative traces of (A) OCR and (B) ECAR from C6 cells acutely incubated with CGP, followed by oligomycin (oligo), 2,4-dinitrophenol (DNP), and rotenone plus antimycin A (Rot/AA) additions, average ± SEM; (C) CGP-induced, (D) proton leak-associated, (E) ATP-linked, and (F) maximal respirations; (G) ECAR variation after ATP synthase inhibition with oligomycin. Paired 2-way ANOVA followed by Holm-Šidak’s post-hoc test, n = 6 independent experiments. (H) lactate secretion measured from C6 cells incubated with CGP or DMSO for 4 h. *P < 0.05, Student’s t test, n = 4 independent experiments, mean and SD (A,B) or mean and paired measurements (C-H).

**Figure 5 – Supplement 1.**
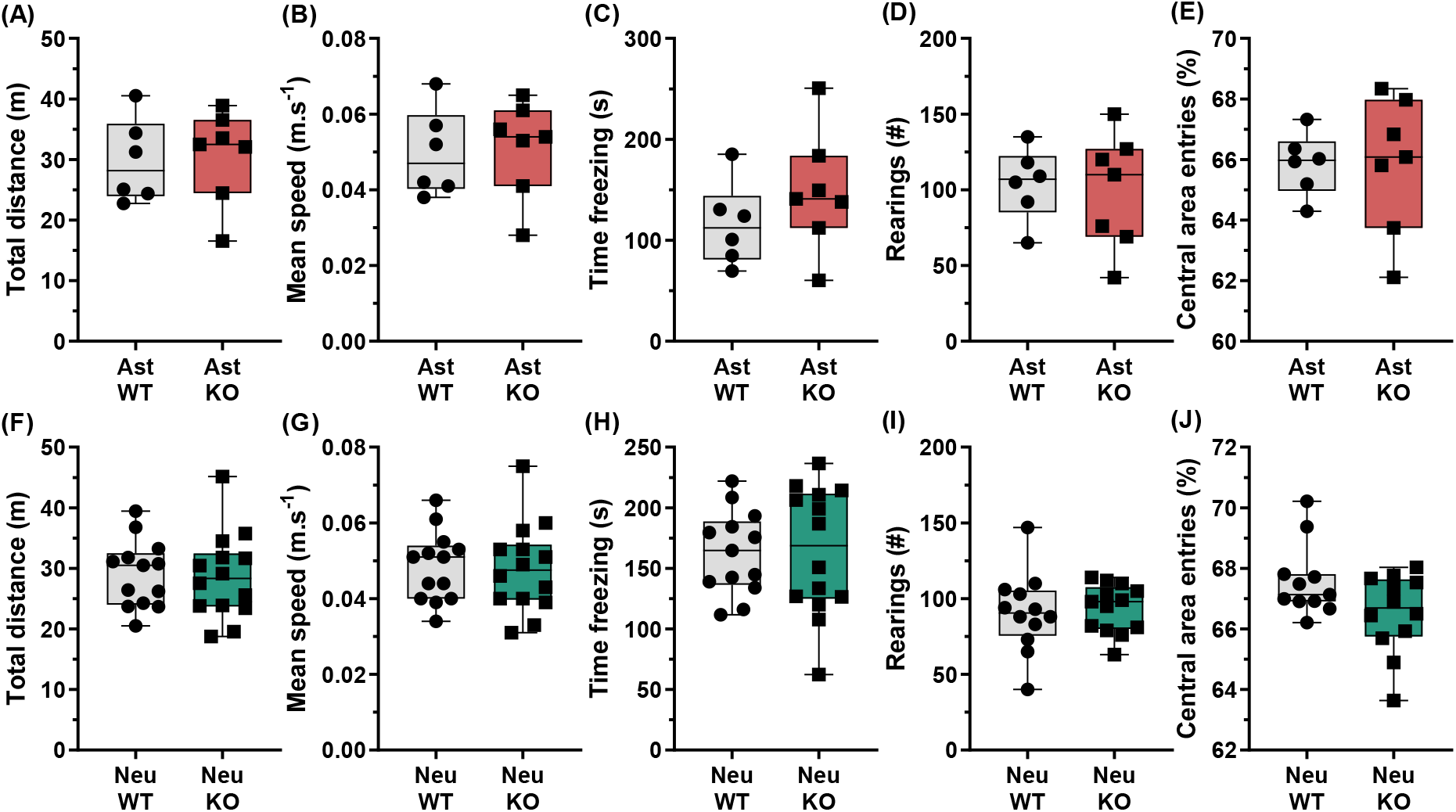
Open Field test supplementary analyses. Supplementary analyses from the Open Field test in mice with hippocampal astrocyte- (A-E) or neuronal-specific (F-J) NCLX deletion: (A,F) total distance; (B,G) mean speed; (C,H) total time in freezing behavior; (D,I) number of rearings; and (E,J) total entries in the central area. Not significant, unpaired Student’s t test (A-C,F-H) or Mann-Whitney test (D,E,I,J), n = 6-14, box indicates upper and lower quartiles and the median (line), and whiskers represent min and max values.

**Figure 5 – Supplement 2. Open Field test – *In vivo* hippocampal astrocyte- and neuronal-specific NCLX deletion representative videos.**

**Figure 5 – Supplement 3.**
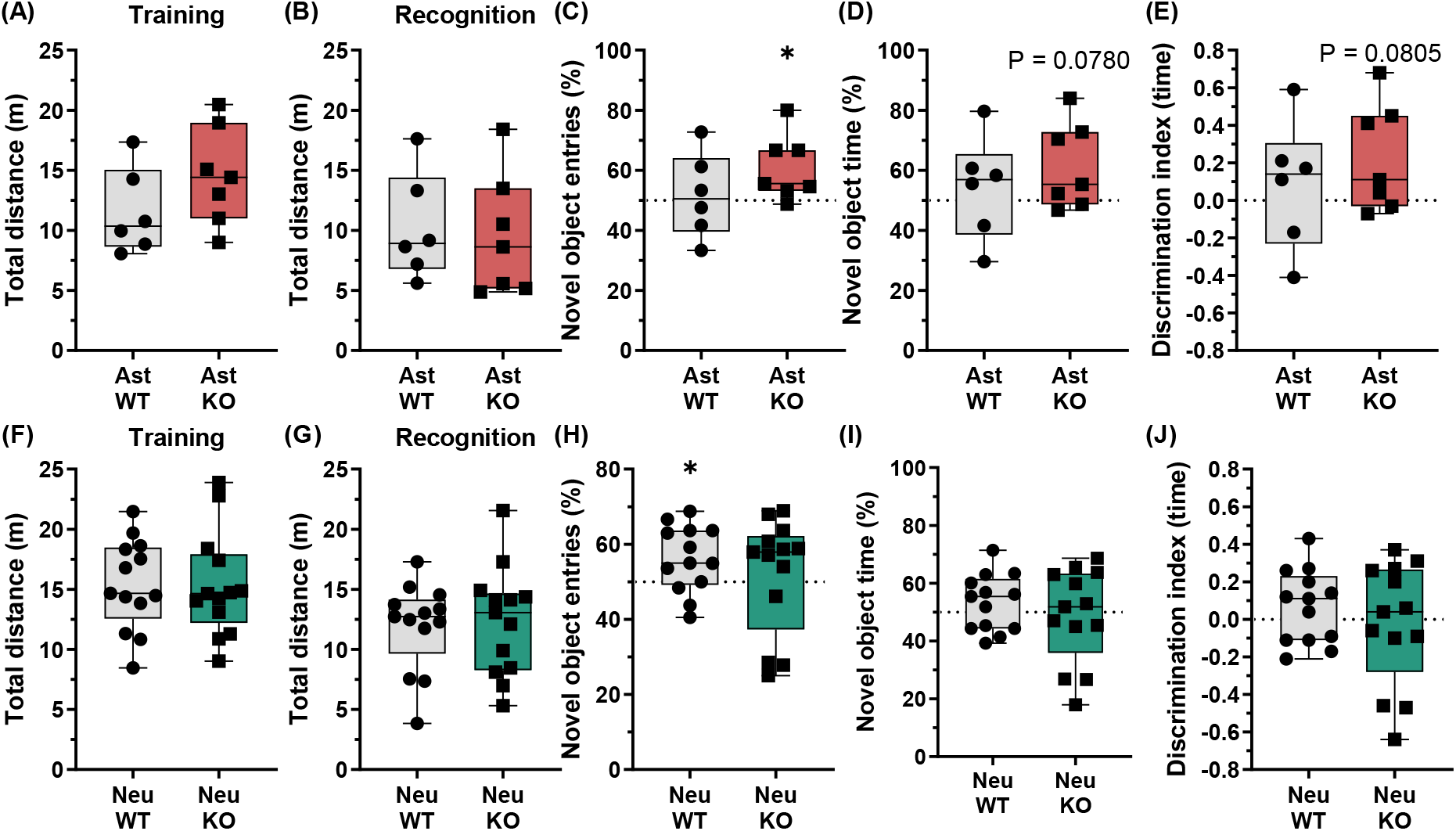
Novel Object Recognition test supplementary analyses. Supplementary analyses from the Novel Object Recognition test in mice with hippocampal astrocyte- (A-E) or neuronal-specific (F-J) NCLX deletion: (A,F) total distance in training and (B,G) recognition assay steps; (C,H) proportion of entries into and (D,I) time within the novel object area; and (E,J) discrimination index calculated from time in the novel and familiar object areas. *P < 0.05, unpaired Student’s t test (A,B,D,E,F,G,H,I) or Mann-Whitney test (C,H), n = 6-14, box indicates upper and lower quartiles and the median (line), and whiskers represent min and max values.

**Figure 5 – Supplement 4. Novel Object Recognition test – *In vivo* hippocampal astrocyte- and neuronal-specific NCLX deletion representative videos.**

**Figure 5 – Supplement 5.**
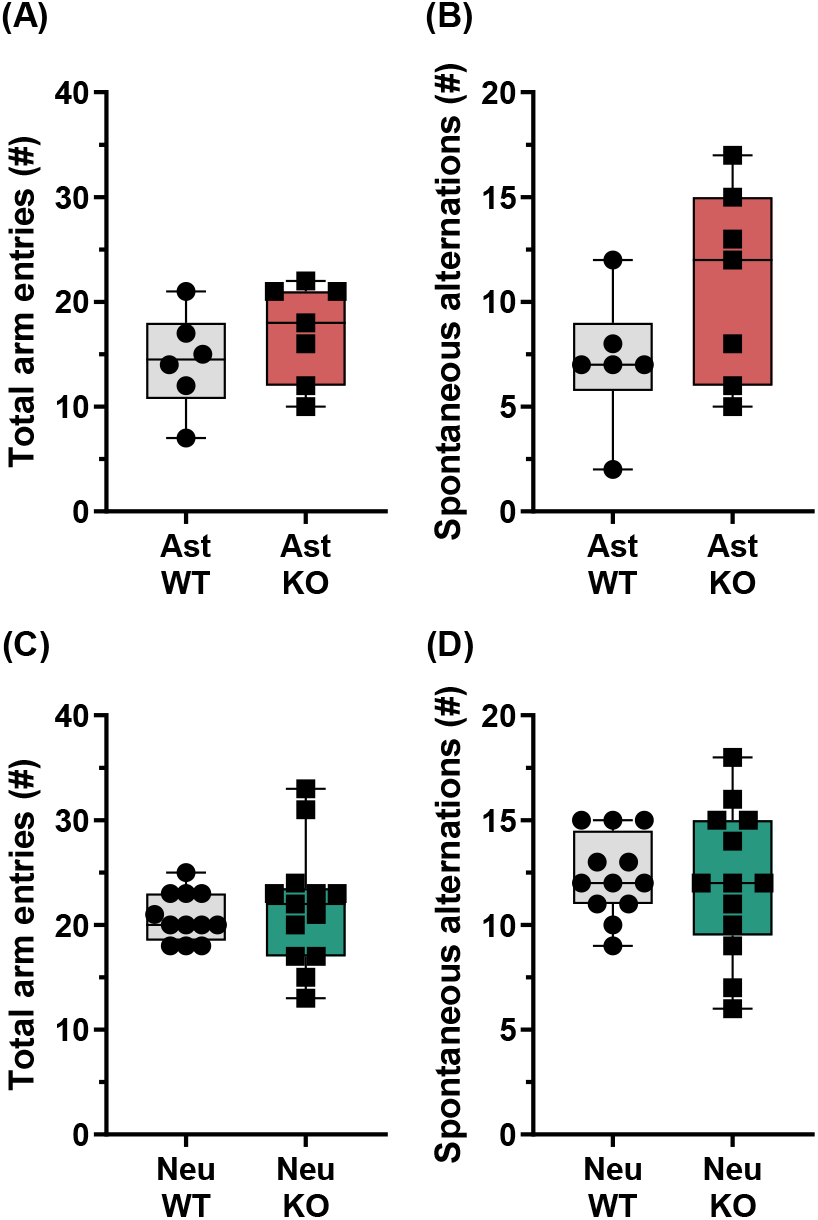
Y-maze test supplementary analyses. Supplementary analyses from the Y-maze test in mice with hippocampal astrocyte- (A,B) or neuronal-specific (C,D) NCLX deletion: (A,C) total number of arm entries; and (B,D) total number of spontaneous alternations. Not significant, Mann-Whitney test, n = 6-14, box indicates upper and lower quartiles and the median (line), and whiskers represent min and max values.

**Figure 5 – Supplement 6. Y-maze test – *In vivo* hippocampal astrocyte- and neuronal-specific NCLX deletion representative videos.**

## Notes

https://osf.io/gjnx5/?view_only=c0db2633132d4dedaa887cc5f9d829a4

### Competing Interest Statement

The authors have declared no competing interest.

